# Identification of Wnt regulated genes that are repressed by, or independent of, β-catenin

**DOI:** 10.1101/2025.04.06.647504

**Authors:** Shiyang Liu, Sara Haghani, Enrico Petretto, Babita Madan, Nathan Harmston, David M Virshup

**Author notes:** Address correspondence to Babita Madan, Nathan Harmston and David M Virshup. Lead author: DMV.

## Abstract

Wnt signaling regulates metazoan development and homeostasis, in part by β-catenin dependent activation and repression of a large number of genes. However, Wnt signaling also regulates genes independent of β-catenin, genes that are less well characterized. In this study, using a pan-Wnt inhibitor we performed a comprehensive transcriptome analysis in a Wnt-addicted orthotopic cancer model to delineate the β-catenin-dependent and independent arms of Wnt signaling. We find that while a large percentage of Wnt-regulated genes are regulated by β-catenin, ten percent of these genes are regulated independent of β-catenin. Interestingly, a large proportion of these β-catenin independent genes are Wnt-repressed. Among the β-catenin dependent genes, more than half are repressed by β-catenin. We used this dataset to investigate the mechanisms by which Wnt/β-catenin signaling represses gene expression, revealing the role of a cis-regulatory motif, the negative regulatory element (NRE). The NRE motif is enriched in the promoters of β-catenin repressed genes and is required for their repression. This provides a comprehensive analysis of the β-catenin independent arm of the Wnt signaling pathway in a cancer model and illustrates how a cis-regulatory grammar can determine Wnt-dependent gene activation versus repression.

## Introduction

Wnt signaling is an evolutionarily conserved pathway involved in diverse processes including development, homeostasis and tissue regeneration (*1*). Dysregulation of this pathway is implicated in myriad diseases including cancer, cardiometabolic disorders and neurodegeneration (*2*, *3*). Signaling is initiated by the binding of Wnt ligands to Frizzleds and other integral membrane co-receptors, which subsequently leads to the activation of distinct downstream signaling pathways. These pathways can operate either through, or independently of, β-catenin, and can either activate or repress specific target genes (*4–7*).

In the β-catenin-dependent pathway, also known as the canonical pathway, the binding of Wnt ligands to their cognate Frizzled receptors results in the stabilization of β-catenin. This stabilized β-catenin translocates into the nucleus and binds to members of the TCF/LEF family of transcription factors to regulate Wnt-target gene expression in a context-dependent manner (*2*, *8*). In this pathway, the β-catenin/TCF complex binds to DNA through the TCF binding motif, also known as the Wnt-responsive element (WRE) (*9*). The β-catenin independent signaling pathways, also known as non-canonical signaling, includes the Wnt/Calcium, Wnt/JNK, Wnt/STOP and planar cell polarity signaling, many of which involve non-canonical Wnts (e.g., WNT5A) and alternative receptors such as ROR. These non-canonical pathways have been implicated in several key cellular processes including migration, planar cell polarity and adhesion that are essential for development and tumorigenesis (*10*, *11*). In contrast to canonical Wnt signaling, our knowledge of the signaling mechanisms and potential target genes in the β-catenin independent arm of the Wnt signaling pathway is less well developed.

The high frequency of mutations leading to aberrant Wnt signaling in multiple tumor types and the changes in transcriptional and cellular states driven by these mutations (*12–15*) has led to the development of pharmacological approaches to inhibit the pathway (*16–18*). One approach is to target Wnt secretion. The post-translational addition of a palmitoleate group to Wnt proteins is necessary for the secretion of all Wnts and is also required for binding to their cognate Frizzled receptor (*19–21*). This palmitoleation is catalyzed by the acyltransferase Porcupine (PORCN)(*22*). Treatment with small molecule inhibitors of PORCN such as ETC-159 and LGK-974 prevents Wnt palmitoleation and the subsequent inhibition of both the β-catenin-dependent and independent branches of Wnt signaling (*23*).

Wnt signaling is generally thought of as a pathway for driving the expression of genes, with most of the well characterized Wnt target genes being Wnt-activated, e.g. *AXIN2, MYC* and *Cyclin D1* (*24–26*). In contrast, only a limited number of genes repressed by Wnt signaling have been well characterized, *e.g. Mmp7* in mice (*27*) and *dpp*, *tig*, d*ugt36Bc* in *Drosophila* (*28–30*) and *BGLAP*, *CDH1* and *CDKN2A* (*31*, *32*) in mammalian cells. In our studies investigating the transcriptional response to a pan-Wnt inhibitor in multiple models of Wnt-driven pancreatic and colorectal cancers (*7*, *18*), we found that Wnt signaling induces the expression of many genes, *i.e. Wnt-activated*, but that a comparable number of genes were upregulated following Wnt inhibition, hence they were *Wnt-repressed*. Further investigation identified that a subset of these *Wnt-repressed* genes were dependent on the inhibition of MAPK signaling by Wnt/β-catenin signaling (*6*, *33*). However, the mechanisms and transcriptional elements involved in Wnt signalling-mediated gene repression and the role of β-catenin in this repression is not well understood.

In this study, we performed a comprehensive transcriptome analysis to delineate the β-catenin dependent and independent arms of Wnt signaling. We used a sensitive orthotopic xenograft model of Wnt-driven pancreatic cancer and compared the transcriptional response to a Wnt-secretion inhibitor, ETC-159 in pancreatic tumors without or with ectopically expressed stabilized β-catenin. This analysis revealed that ∼90% of Wnt-dependent genes are regulated by β-catenin, while only ∼10% are regulated independently of β-catenin. The same dataset was interrogated to better understand how Wnt/β-catenin signaling can repress gene expression. This analysis identified an enrichment of a specific negative regulatory element (NRE) in the promoters of the β-catenin-dependent Wnt-repressed (*34*). Our data supports the role of the NRE as an important cis-regulatory motif required for the regulation of β-catenin dependent genes in human cells. This suggests the existence of a cis-regulatory grammar which may be responsible for determining whether a target gene will be repressed or activated by Wnt signaling.

## Results

### Identification of β-catenin-dependent and -independent Wnt target genes

HPAF-II cells have an inactivating mutation in RNF43 that drives high autocrine Wnt signaling, making them Wnt-addicted and sensitive to treatment with PORCN inhibitors such as ETC-159 (*18*, *35*). In this context, PORCN inhibition leads to the ablation of both β-catenin dependent and independent Wnt signaling. To dissect the differences between these two branches of the Wnt signaling pathway, we generated HPAF-II cells with constitutively active β-catenin dependent signaling. This was accomplished by stably transducing HPAF-II cells with a plasmid expressing β-catenin with four phosphorylation sites (S33, S37, T41, S45) mutated to alanine, referred to here as β-cat4A (*36*). Phosphorylation at these sites by CK1α and GSK3 is required to target β-catenin for proteasomal degradation. As such, treatment of β-cat4A cells with ETC-159 will only affect the expression of Wnt-dependent but β-catenin independent target genes, while genes under the control of β-catenin will be unaffected (**Figure S1A**).

Clones stably expressing β-cat4A were established by single-cell cloning and clones with near-physiological expression levels were selected (**Figure 1A**). To assess the ligand-independent activation of Wnt/β-catenin signaling in these cells we measured their sensitivity to the pan-Wnt inhibitor ETC-159. Confirming the feasibility of this approach, in parental HPAF-II (WT) cells, ETC-159 treatment led to a dose-dependent decrease in the expression of the well-characterized Wnt/β-catenin target gene *AXIN2*, while in β-cat4A cells *AXIN2* expression increased at baseline and was not downregulated by Wnt inhibition (**Figure 1B**). Moreover, in a soft agar assay there was only a slight decrease in the β-cat4A colonies even in the presence of 100 nM (∼30x the IC50) ETC-159 (**Figure 1C**). This demonstrates that the growth of β-cat4A cells in vitro largely requires Wnts to activate β-catenin signaling.

**Figure 1.**
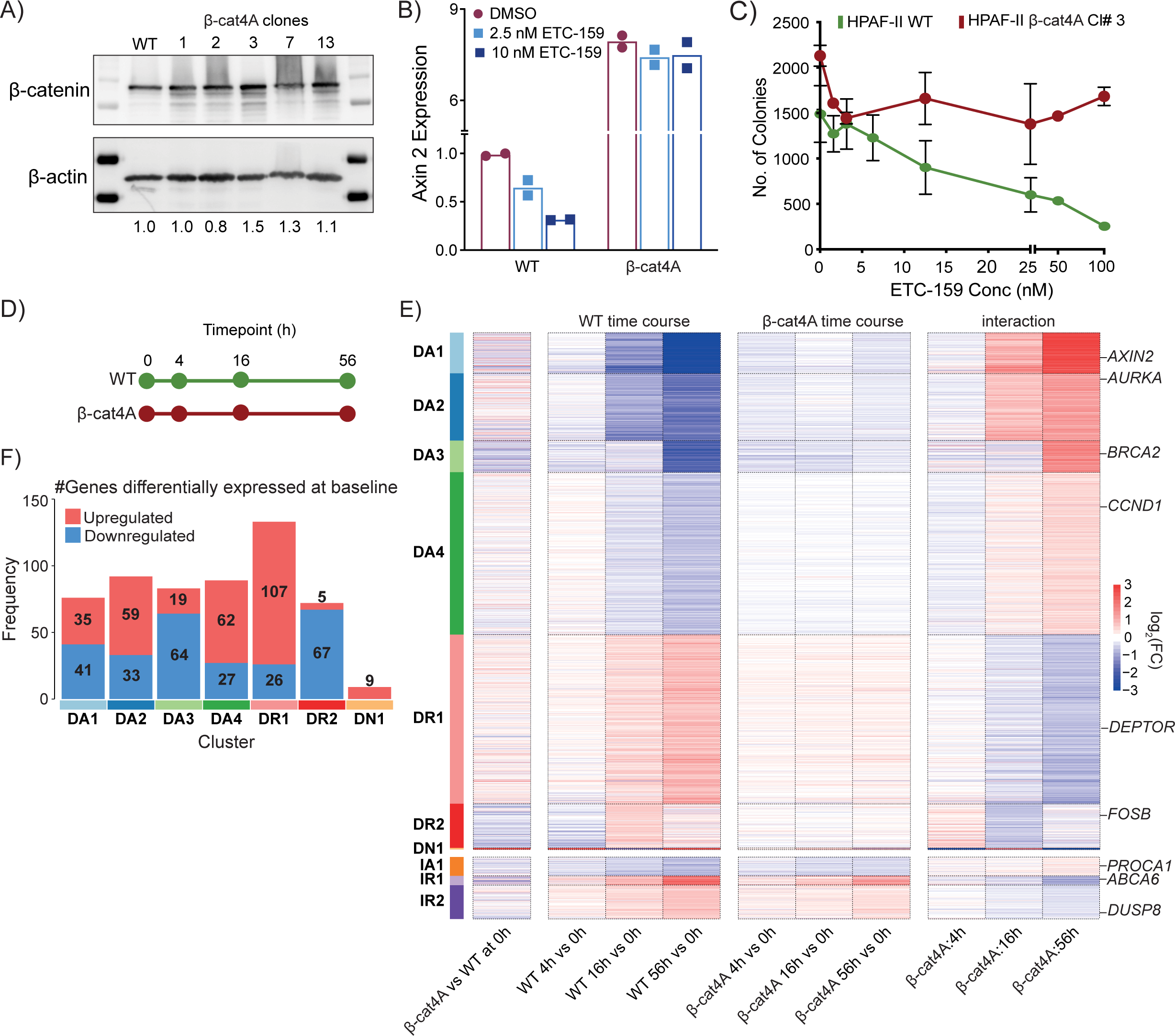
Identification and classification of genes regulated by distinct branches of the Wnt signaling pathway. A) Near-physiologic expression of β-cat4A in HPAF-II cells. WT = parental cell line. The numbers under the blot indicate the normalized ratio of β-catenin to β-actin. B) β-cat4A abrogates the loss of *AXIN2* expression as assessed by RT-qPCR following treatment of cells with indicated concentration of the PORCN inhibitor ETC-159. C) β-cat4A expressing cells are able to form colonies despite PORCN inhibition, while WT cells show a dose-dependent loss of the ability to form colonies (p=0.0034). D) WT and β-cat4A tumors were harvested for RNA-seq at four distinct timepoints following treatment with ETC-159 (37.5 mg/kg every 12 hours by gavage). E) Differential expression analysis identified 3,346 genes that sorted into seven β-catenin dependent clusters and three β-catenin independent clusters of genes (see Methods). Heatmap shows log2 fold change for multiple comparisons of interest. β-cat4A vs WT at 0 h shows the differences in expression between the two conditions at 0 h. Changes in gene expression over time (compared to 0 h) are shown for tumors generated from WT HPAF-II cells (WT timecourse) and mutant β-catenin cells (β-cat4A timecourse). Also displayed are interaction terms representing the difference between these values after controlling for differences in baseline expression. DA is dependent, activated, DR is dependent, repressed, DN is dependent noise, IA in independent activated, IR is independent repressed. F) 554 genes were identified as significantly differentially expressed at baseline (WT 0 h vs β-cat4A 0 h), distributed across each of the β-catenin dependent clusters.

To identify β-catenin dependent and independent genes in a more physiological setting, we used an orthotopic xenograft model, where the transcriptional response to Wnt inhibition is significantly more robust than it is *in vitro* or in flank xenografts (*7*). WT and β-cat4A HPAF-II cells were injected into the mouse pancreas. Following tumor establishment pan-Wnt inhibition was achieved with ETC-159 (37.5 mg/kg b.i.d. orally) treatment (**Figure 1D**). Gene expression changes were assessed at 4, 16, and 56 h of treatment by RNAseq. Genes were classified as β-catenin dependent or independent based on their transcriptional response to PORCN inhibition in the presence or absence of stabilized β-catenin (**Table S1**). β-catenin dependent genes were defined as those that were differentially expressed over time in the WT condition (FDR<0.1) and responded differently in WT versus β-cat4A tumors (interaction test, FDR<0.1). These criteria resulted in 2988 genes being classified as transcriptional targets of β-catenin dependent signaling. β-catenin independent genes, likely regulated by Wnt-dependent non-canonical pathways, were those that were differentially expressed over time after Wnt inhibition in both β-catenin WT (FDR<0.1) and β-cat4A (FDR<0.1) conditions and whose response to ETC-159 treatment did not significantly differ between conditions (interaction test, FDR>0.1). Using these criteria, 358 genes (∼10% of the total number of Wnt-regulated genes) were classified as β-catenin-independent.

To better understand the changes in gene expression, these two sets of genes were clustered based on their temporal response to Wnt inhibition. This identified seven clusters of β-catenin-dependent genes, and three clusters of β-catenin independent genes (**Figure 1E**). DA1-4 (DA = **<u>D</u>**ependent & **<u>A</u>**ctivated) were classified as β-catenin dependent and Wnt-activated, as their expression decreased in response to ETC-159, while DR1-2 (DA = **<u>D</u>**ependent & **<u>R</u>**epressed) were classified as β-catenin dependent and Wnt-repressed, as their expression increased in response to ETC-159 treatment. DN1 (**<u>D</u>**ependent **<u>N</u>**oise) consisted of only nine genes, likely due to clustering artefact. DA1 and DA3 contain most of the well-known direct Wnt-target genes (e.g. *AXIN2, NOTUM, RNF43, MYC, NKD1, BMP4*).

As a proof of concept, we examined *AXIN2*, a well-known direct Wnt-regulated β-catenin target gene. As expected, our analysis classified *AXIN2* as a β-catenin dependent Wnt-activated gene (DA1). In orthotopic HPAF-II tumors, inhibition of Wnt/β-catenin signaling by ETC-159 led to a dramatic downregulation of *AXIN2* expression (FDR=2.58×10^-132^) (**Figure 2A).** However, in the presence of mutant β-catenin, baseline *AXIN2* expression increased, and Wnt inhibition had no further effect (FDR=0.49). When comparing the expression changes following Wnt inhibition, a significant interaction was observed (interaction test, FDR=196×10^-58^), indicating a differential response to Wnt inhibition over time depending on the status of β-catenin. Conversely, *DEPTOR* was identified as a β-catenin dependent, Wnt-repressed gene (DR1). It was significantly upregulated in WT tumors following Wnt inhibition (FDR=7.68×10^-11^), but did not respond to ETC-159 treatment in the presence of mutant β-catenin (FDR=0.52), and these responses were significantly different between the two conditions (interaction test, FDR=0.06) (**Figure 2B**).

**Figure 2.**
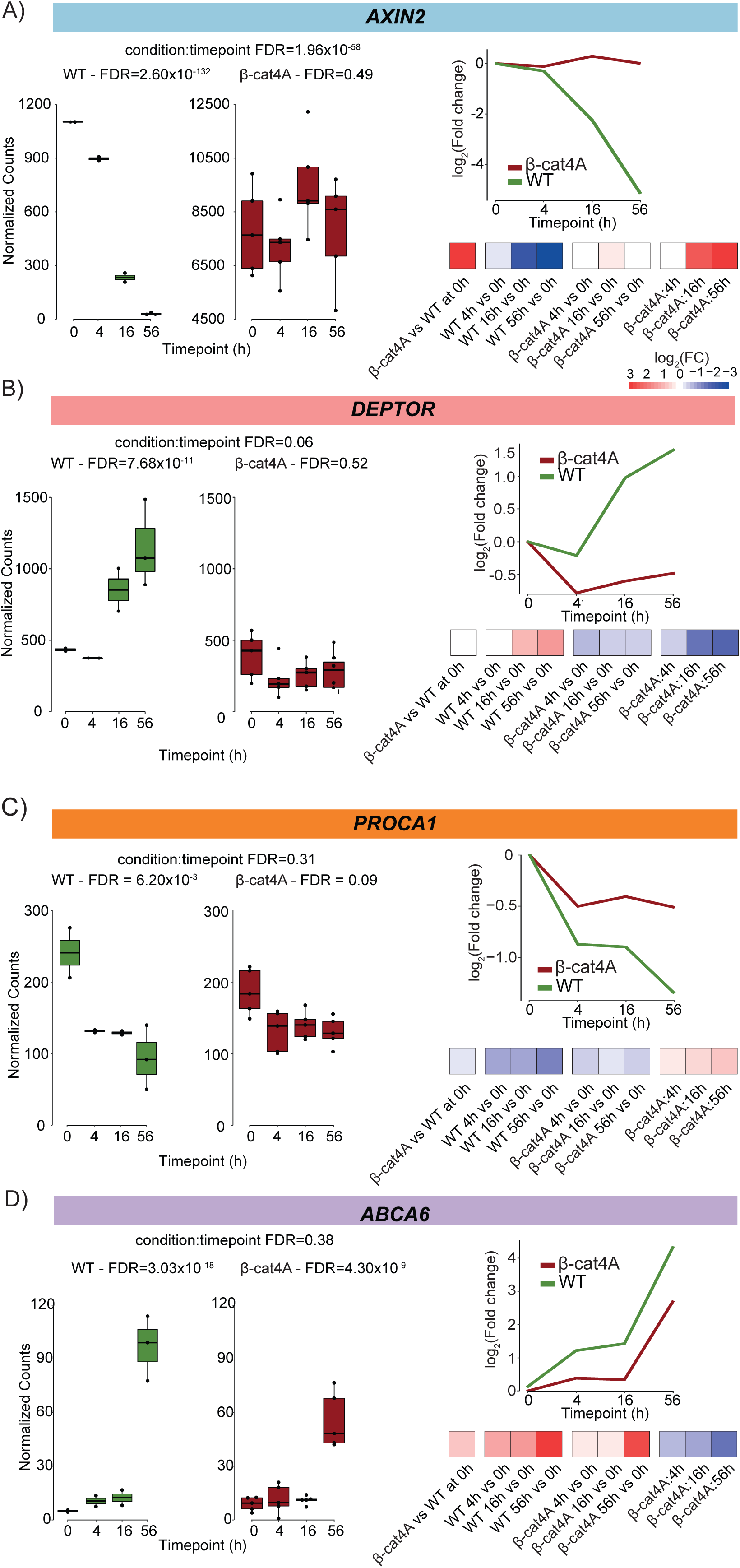
Identification of β-catenin dependent and -independent genes. Examples of Wnt target genes identified in this analysis. For each gene, the expression (normalized counts) of the gene is shown, as are the fold changes and how these changes correspond to the heatmap representation used in Figure 1E. A) *AXIN2* is a β-catenin dependent Wnt-activated gene, in cluster DA1. B) *DEPTOR* is a β-catenin dependent Wnt-repressed gene (DR1). C) *PROCA1* is a β-catenin independent Wnt-activated gene (IA1). D) *ABCA6* is a β-catenin independent Wnt-repressed gene (IR1).

Three clusters were identified in the 358 β-catenin independent genes. Cluster IA1 (**<u>I</u>**ndependent & **<u>A</u>**ctivated) was Wnt-activated, while both IR1 and IR2 were Wnt-repressed. *PROCA1* (IA1) and *ABCA6* (IR1) are examples of β-catenin independent genes (**Figures 2C** and **2D**) that responded to PORCN inhibition in both the WT and β-cat4A tumors with no significant difference in their response, regardless of the β-catenin protein abundance. Genes in these clusters are presumably regulated by non-canonical Wnt signaling pathways, either directly or indirectly.

As an independent validation that our approach identified *bona fide* β-catenin independent targets, we treated the β-cat4A and WT cells with a tankyrase inhibitor, G007LK. G007LK treatment alters β-catenin abundance, impacting the expression of β-catenin target genes without affecting β-catenin independent targets (*37*). Similar to the effect of the PORCN inhibitor, G007LK treatment reduced *AXIN2* expression (**Figure S1B)**, but did not reduce the expression of the β-catenin independent genes *KRT19* and *DUSP5* (**Figure S1C-D)**.

HPAF-II tumors are dependent on continuous Wnt signaling, so it was of interest to determine if a further increase in β-catenin caused by ectopic expression of β-cat4A would further change gene expression, or if β-catenin was near-saturating in this system. Of the 2988 β-catenin-dependent Wnt-regulated genes in the HPAF-II orthotopic tumors, only ∼10% (296) had significantly higher expression in the tumors with stabilized β-catenin, while 8.6% (258) had a significantly lower expression compared to the WT tumors (**Figure 1E**, column 1). Significant differences were defined as absolute fold-change > 1.5, FDR < 0.1 from their baseline expression, *i.e.* between β-cat4A and WT (**Figure 1F**). As expected, no genes identified as β-catenin-independent were differentially expressed at baseline. This data suggests that this Wnt-addicted cancer has reached close to maximal β-catenin activation.

In summary, transcriptional profiling of our orthotopic *in vivo* model identified a robust set of β-catenin-dependent and -independent target genes, with the β-catenin independent, non-canonical genes accounting for only 10% of the differentially expressed genes in this model. This great difference between the number of β-catenin dependent and β-catenin independent genes indicates that, as least in this context, WNT signaling regulates gene expression predominantly through β-catenin, and/or that non-canonical Wnt signaling may predominantly be acting via non-transcriptional mechanisms.

### β-catenin-dependent and -independent Wnt target genes associate with distinct biological pathways

Functional enrichment analysis was performed to characterize the clusters of β-catenin dependent and independent genes (**Figure 3A, Table S2**). The clusters of Wnt-activated β-catenin dependent genes (DA1-4) were enriched for processes and pathways including Wnt signaling, ribosome biogenesis, DNA replication, splicing and DNA repair, while the Wnt-repressed β-catenin dependent genes (DR1-2) were enriched for protein transport and EGF signaling pathways. This corroborates an extensive literature on Wnt target genes and our previous findings dissecting the effects of inhibiting the Wnt signaling pathway in RNF43 mutant pancreatic cancer (*6*, *7*, *33*, *38*, *39*) and confirms that these processes are regulated downstream of β-catenin.

**Figure 3:**
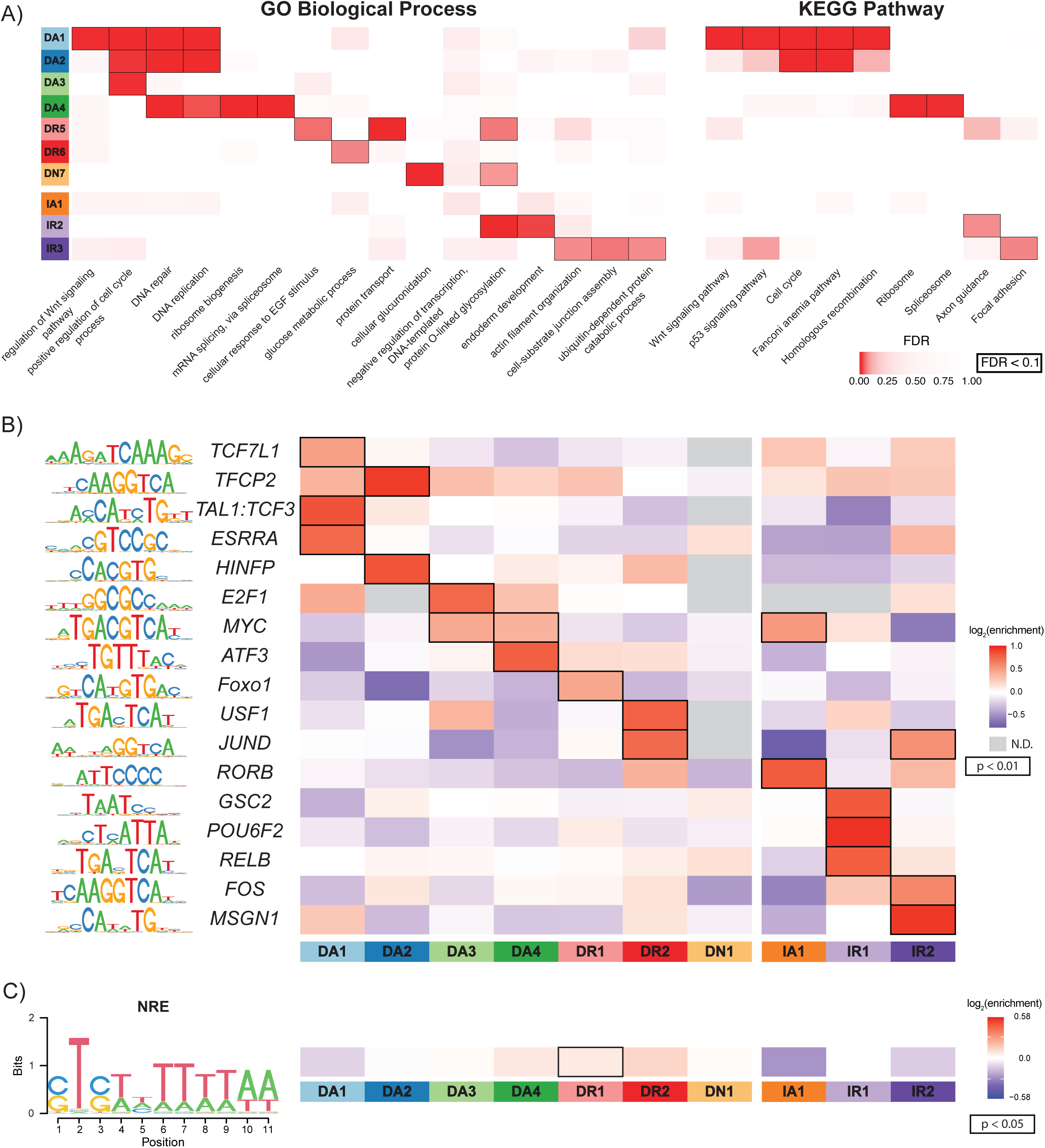
β-catenin independent and dependent clusters are enriched for distinct pathways, processes and TFBS motifs. A) Enrichments for GO biological processes and KEGG pathways for each of the clusters of β-catenin dependent and independent genes. B) TFBS motifs enriched in the promoters of β-catenin dependent and independent genes. Enrichment is calculated as observed divided by expected. C) Sequence logo of the NRE motif, modified after (*34*), is found in multiple clusters and significantly enriched in a cluster of β-catenin-dependent Wnt-repressed genes (DR1).

In contrast, the clusters of β-catenin independent Wnt-activated genes in IA1 showed no significant pathway enrichment. However, genes in the IR2 cluster showed enrichment for processes related to endoderm development, protein O-linked glycosylation and axon guidance, and those in IR1 were enriched for actin organization and cell junction assembly. Taken together, this indicates that the β-catenin dependent and independent branches of the Wnt signaling pathways regulate largely distinct downstream signaling pathways and processes, with non-canonical signaling affecting processes related to development and tissue organization.

### β-catenin dependent and independent genes are enriched for distinct transcription factor binding sites

To identify the transcription factors potentially involved in regulating each of the β-catenin dependent and independent clusters we performed Transcription Factor Binding Site (TFBS) enrichment analysis (**Table S2**) on their promoters, examining sequences 2 kb upstream and 500 bp downstream from their respective transcriptional start site (TSS) (Figure 3B).

β-catenin most famously regulates gene expression by binding to and de-repressing members of the LEF/TCF family of transcription factors, thus activating transcription. Consistent with this, DA1, the cluster containing most of the well-known β-catenin dependent genes showed significant enrichment for TCF7L1 motif, also known as the Wnt-response element, or WRE. DA3 and DA4, clusters enriched for cell cycle related genes (Figure 3A) were also enriched for binding sites for E2F1 and MYC and other key mediators of cell proliferation and mitosis. We note that many relevant LEF/TCF binding sites may be present in enhancers rather than promoters, explaining why they are not strongly enriched in all β-catenin dependent Wnt activated gene clusters. On the other hand, β-catenin-dependent repressed clusters were significantly enriched for binding sites for USF1, JUND and FOXO1.

The β-catenin-independent genes were enriched for a distinct set of TFBSs, with the activated genes showing significant enrichment for RORB binding sites. The β-catenin independent repressed genes were notably enriched for motifs bound by homeobox factors including GSC2, POU6F2, and MSGN1. This finding aligns with the known role of non-canonical Wnt signaling in embryonic development. Overall, the set of enriched motifs were distinct for the dependent and independent genes, consistent with our current understanding that they are regulated by distinct signaling pathways.

### A Negative regulatory element (NRE) is enriched in Wnt-dependent gene clusters

An 11-bp sequence, known as the Negative Regulatory Element (NRE) (**Figure 3C**), was previously identified as a motif that modulates the expression of Wnt/β-catenin target genes in *Xenopus laevis*, and was also shown to be functional in mouse embryonic stem cells (*34*). This sequence was shown to recruit both TCF and β-catenin, with the binding of both proteins being necessary to mediate its repressive effects. We investigated whether any of the identified clusters of Wnt-regulated genes were enriched for this motif (**Figure 3C**, Methods). Indeed, we observed an enrichment for a variant of the published NRE element in the Wnt/β-catenin dependent repressed gene clusters with significant enrichment in DR1, suggesting that this motif might be responsible for mediating the expression of a subset of Wnt-repressed target genes.

### The NRE motif is enriched in TCF4 and β-catenin ChIP-seq bound peaks

Given the enrichment for the NRE motif in Wnt/β-catenin-repressed genes, we examined publicly available β-catenin and TCF4 (the protein product of the *TCF7L2* gene) ChIP-seq datasets to identify whether the NRE motif was present in regions bound by either of these proteins (*40*, *41*). Analysis of the TCF4 ChIP-seq data obtained from six cell lines found that both the WRE motif and NRE motif were significantly enriched in the TCF4 peaks in all cell lines (**Figure 4A**, Bonferroni corrected p-value < 0.05). The NRE was also significantly enriched compared to random background sequences in the other seven cell-lines, although the enrichment was consistently lower compared to the enrichment observed for the TCF4-binding WRE (**Figure 4A** and **B**). Analysis of publicly available β-catenin ChIP-seq data generated from DLD1 and SW480 cells also revealed that β-catenin peaks were significantly enriched for both NREs and WREs. In five of the eight cell lines investigated, these two motifs were found to significantly co-occur within peaks more than expected by chance (Fisher’s exact test, Bonferroni corrected p-value < 0.05) (**Figure 4C**); however, no preferred distance between NREs and WREs relative to each other was observed (**Supplemental Figure 2A**). This supports the proposed role of WREs and NREs functioning together to regulate the expression of a target genes in response to Wnt signaling (*34*).

**Figure 4:**
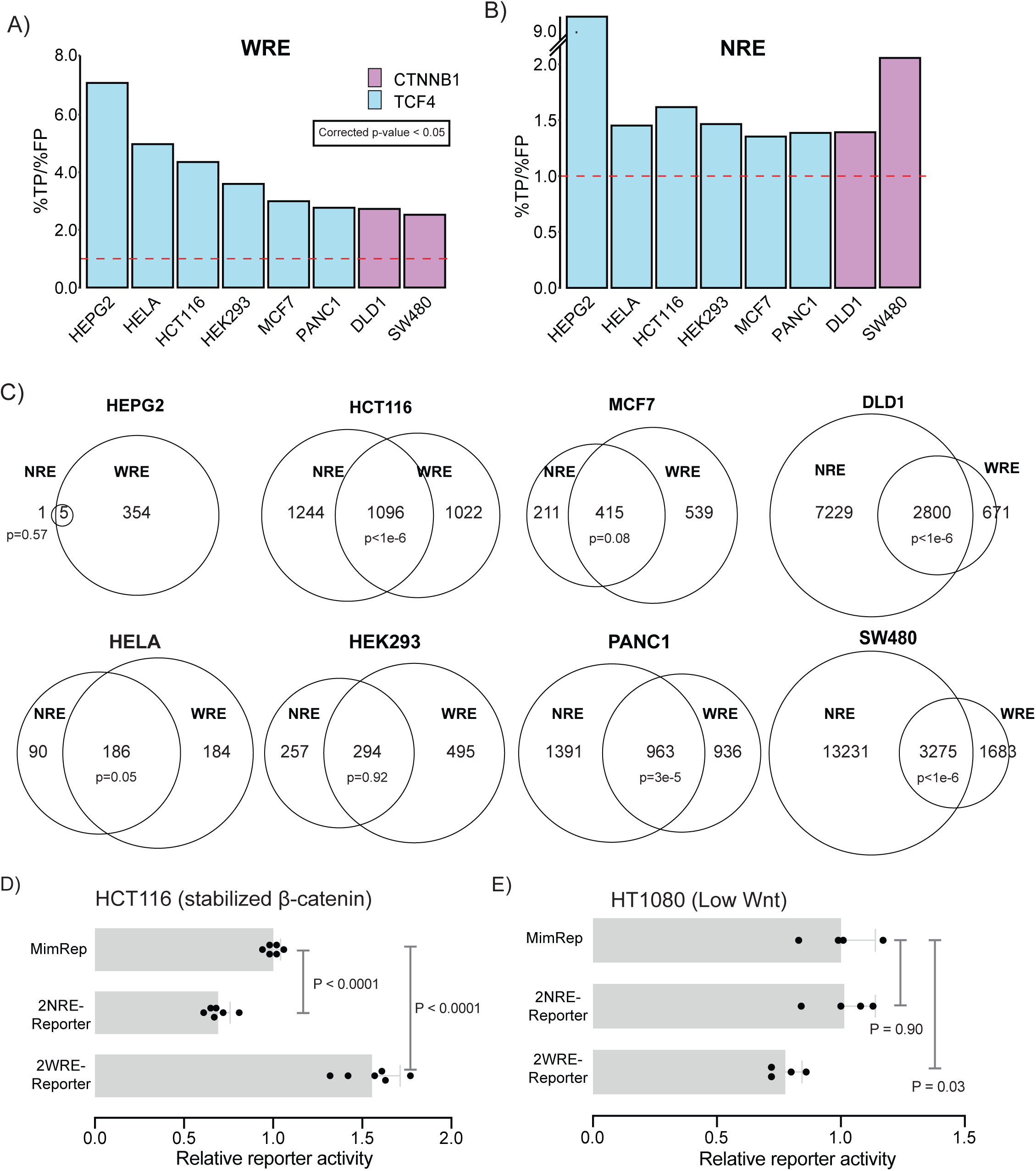
The NRE is capable of repressing reporter activity and is found at binding sites of TCF7L2/β-catenin. A) Enrichment of WREs across multiple TCF7L2/β-catenin ChIP-seq datasets. %TP/%FP represents the ratio of the percentage of peaks identified as containing a motif (%TP) versus percentage of background sequences containing a motif (%FP). B) Enrichment of NREs across the same datasets. While HEPG2 shows strong enrichment for NREs and WREs, it only has a small number of peaks in the dataset. C) WREs and NREs co-occur more often than predicted by chance across TCF7L2/β-catenin peaks identified in multiple cell lines. Venn diagrams show the number of WRE and NRE motifs identified and the overlap between them; significance was calculated using Fishers Exact test. D) Addition of NREs represses, while WREs activates a minimal reporter (MimRep) in HCT116 cells driven by stabilized β-catenin. E) The same reporters are not Wnt-regulated in HT1080 Wnt-low cells.

### The negative regulatory element is sufficient to mediate Wnt signaling induced transcriptional repression in human cells

To functionally assess the role of the NRE in repressing genes in a β-catenin-dependent manner we created three synthetic reporters: i) a minimal reporter (MimRep) that does not respond to Wnt signaling (**Supplementary Figures 2B**), ii) a 2NRE-reporter containing two 11 bp NRE sites, and iii) a 2WRE-reporter with two WRE sites placed in front of the minimal reporter. In human colorectal cancer HCT116 cells that have hyperactivated Wnt signaling due to a S45del mutation in β-catenin, as expected the 2WRE-reporter showed significantly increased activity, while the 2NRE-reporter activity was repressed compared to the minimal reporter (**Figure 4D**). In contrast, in human fibrosarcoma HT1080 cells with low basal Wnt activity, both the 2NRE-reporter and minimal reporter had similar transcriptional activities, while the 2WRE-reporter showed reduced transcriptional activity (**Figure 4E**). This reduction could potentially be due to the interaction between TCF/LEF and Groucho in the “Wnt-off” condition, which is shown to mediate transcriptional repression of the Wnt activated genes via the WRE (*42*, *43*). These findings suggest that the NRE motif is sufficient to repress reporter activity in a β-catenin-dependent manner in human cancer cell lines.

### NREs are necessary for the repression of the Wnt-repressed/β-catenin-dependent lncRNA *ABHD11-AS1*

As the NRE motif was found to functionally repress reporter activity in HCT116 cells and was enriched in multiple TCF4/β-catenin ChIP-seq datasets, we hypothesized that it may play an important role in regulating a subset of Wnt-repressed/β-catenin-dependent genes in human cancer cells. We previously identified *ABHD11-AS1* as a Wnt-repressed long non-coding RNA (lncRNA) in an orthotopic model of Wnt-addicted pancreatic adenocarcinoma (**Figure 5A**). CRISPRi mediated knockdown of *ABHD11-AS1* led to an increase in the growth of HPAF-II derived subcutaneous tumors, supporting its role as a tumor suppressor *in vivo* (Liu et al., 2020). To test if *ABHD11-AS1* expression is β-catenin-dependent, we treated HT1080 cells with the GSK3 inhibitor BIO that regulates canonical Wnt signaling by stabilizing β-catenin (*44*). Treatment of HT1080 cells with BIO led to a significant decrease in *ABHD11-AS1* expression (**Figure 5C**). In both HPAF-II and HCT116 cells that have hyperactivated Wnt signaling due to mutations in *RNF43* and *CTNNB1* respectively, knocking down *CTNNB1* using either CRISPRi in HPAF-II cells (**Figure 5D**), or siRNA in HCT116 cells (**Figure 5E**) led to a significant increase in *ABHD11-AS1* expression. Furthermore, ETC-159 treatment of HPAF-II cells, inhibiting Wnt secretion, increased *ABHD11-AS1* expression, but this effect was blocked in the presence of stabilized β-catenin (**Figure 5F**). Taken together, these results indicate that *ABHD11-AS1* lncRNA is repressed by β-catenin across multiple cancer cell lines.

**Figure 5:**
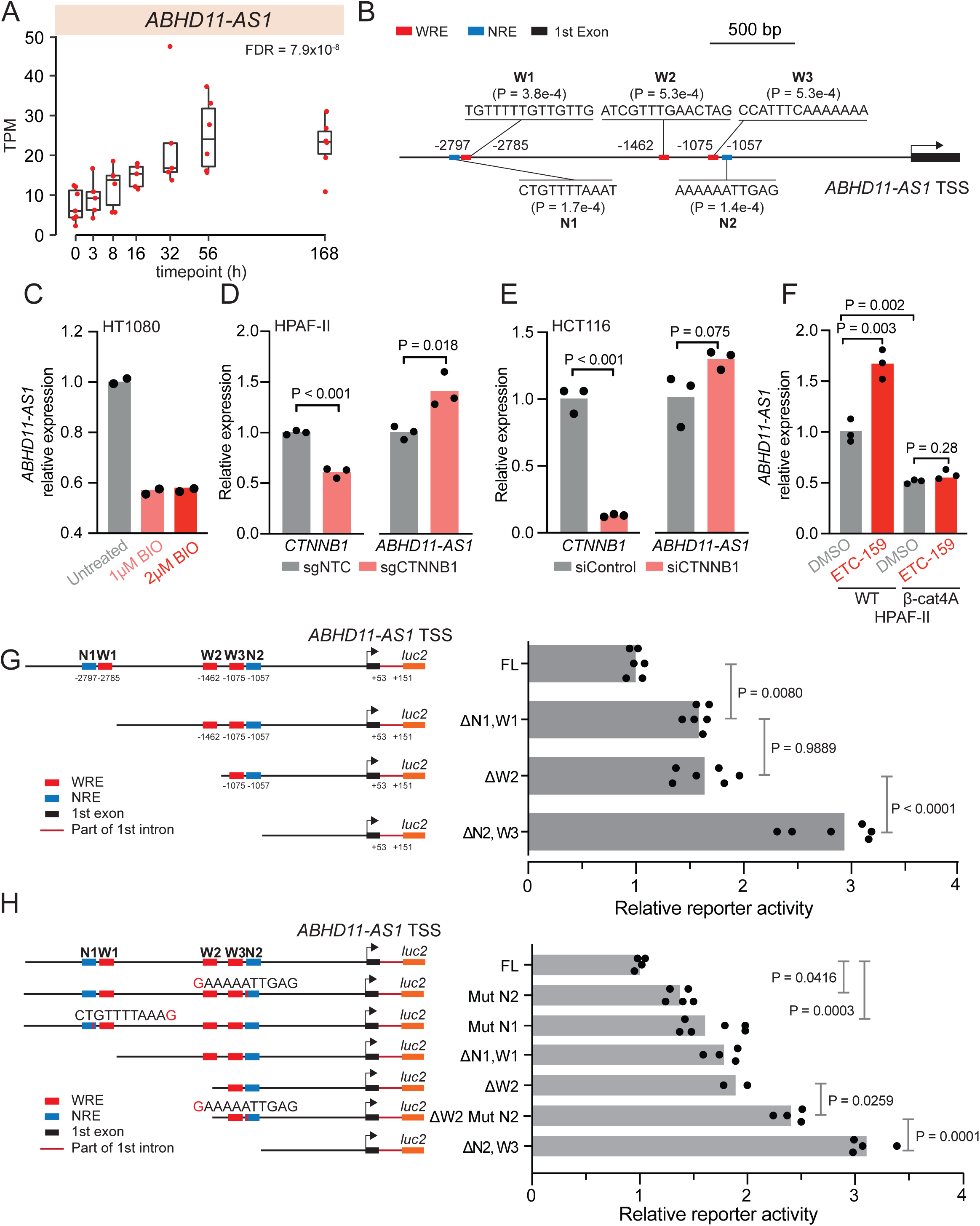
NRE is necessary for the repression of the Wnt-repressed/β-catenin-dependent lncRNA ABHD11-AS1. A) *ABHD11-AS1* is a Wnt-repressed gene in orthotopic model of HPAF-II Wnt-addicted cancer where its expression increases following PORCN inhibition (data from (*7*, *66*) B) Positions of putative WREs and NREs in the promoter of *ABHD11-AS1* C) *ABHD11-AS1* expression is repressed when β-catenin signaling is activated by inhibition of GSK3 with BIO. D) Inhibiting β-catenin using CRISPR or E) siRNA leads to an increase in expression of *ABHD11-AS1* in cultured HPAF-II and HCT116 cells respectively. F) Expression of *ABHD11-AS1* increases in WT but not β-cat4A HPAF-II cells following Wnt inhibition with ETC-159. G) Transcriptional activity of *ABHD11-AS1* reporter construct in HCT116 cells following serial deletion of sequences containing WREs and NREs H) Transcriptional activity of *ABHD11-AS1* reporter construct in HCT116 cells following mutation of the two NRE sites.

We then examined the *ABHD11-AS1* for NREs and WREs. Analysis of the promoter region of *ABHD11-AS1* (3.4 kb upstream and 200 bp downstream of its TSS to the 1st intron) identified two candidate NREs (P < 2×10^-4^), located at 2797 and 1057 bp upstream of the TSS (denoted N1 and N2 respectively), and three candidate WREs (P < 6×10^-4^), located 2785, 1462 and 1075 bp upstream of the TSS (**Figure 5B**). To elucidate the role of NREs in the regulation of *ABHD11-AS1*, we cloned its promoter region, from 3327 bp upstream to 151 bp downstream of its TSS, into a luciferase reporter pGL4.20. We then systematically deleted regions of the promoter to remove NREs and measured the resulting reporter activity in HCT116 cells. Deleting a region containing N1 (ΔN1, W1), led to a 1.6-fold increase in reporter activity compared to the full length (FL) construct (**Figure 5G**), suggesting that NRE (N1) represses *ABHD11-AS1* expression. Further deletion of a 1265 bp fragment (ΔW2), with no identifiable NREs, did not change the transcriptional activity (**Figure 5G**). However, deleting a 400 bp DNA fragment containing N2 (ΔN2, W3), led to a 2.9-fold increase in the reporter activity (**Figure 5G**). This suggests that NREs are required for repressing the expression of *ABHD11-AS1*.

Large deletions of the promoter fragment could potentially lead to the loss of additional functional elements besides the NREs. Therefore, to specifically investigate the effect of NREs on the *ABHD11-AS1* regulation, we performed a series of mutagenesis experiments (**Figure 5H**). It has been shown that mutating the 11th base of the NRE motif can reduce its suppressive function (*34*). We therefore mutated the last base of each of the two NREs (Mut N1, Mut N2) in the 3478-bp promoter fragment. Each of these mutants significantly enhanced reporter activity to a level comparable to that observed following deletion of N1 (ΔN1) harboring regions. However, the reporter activity of the N2 mutant was further increased by deletion of both the NRE and WRE elements (**Figure 5H**, compare ΔW2 Mut N2 with ΔN2, W3), suggesting that a single bp change does not completely abrogate the NRE function, and/or that the interaction with the WRE element is required for the repression. Taken together, these data show that perturbation of NREs in the *ABHD11-AS1* promoter leads to its activation, confirming that NREs are functional motifs that are capable of repressing gene expression in a Wnt-dependent manner in human cancer cells.

### NRE modulates the expression of Wnt-activated/β-catenin-dependent gene *AXIN2*

Kim *et al.* suggested that β-catenin binds to both NREs as well as WREs to modulate the expression of the Wnt-activated genes *siamois* in *Xenopus* and *Brachyury* in mESCs (*34*). To test the hypothesis that NREs can modulate Wnt-activated/β-catenin-dependent genes, we examined *AXIN2,* a well-established β-catenin dependent Wnt-activated gene (**Figure 6A**). Scanning the human *AXIN2* promoter region (from 3.5 kb upstream to 1 kb downstream of its TSS), we identified seven WREs (P < 1.5×10^-4^) (**Figure 6B**) with five located within the first intron and two within the 1 kb of its TSS. In addition, we also identified one NRE (P < 1.5e-04) located 1329 bp upstream from the TSS (Figure 6B).

**Figure 6:**
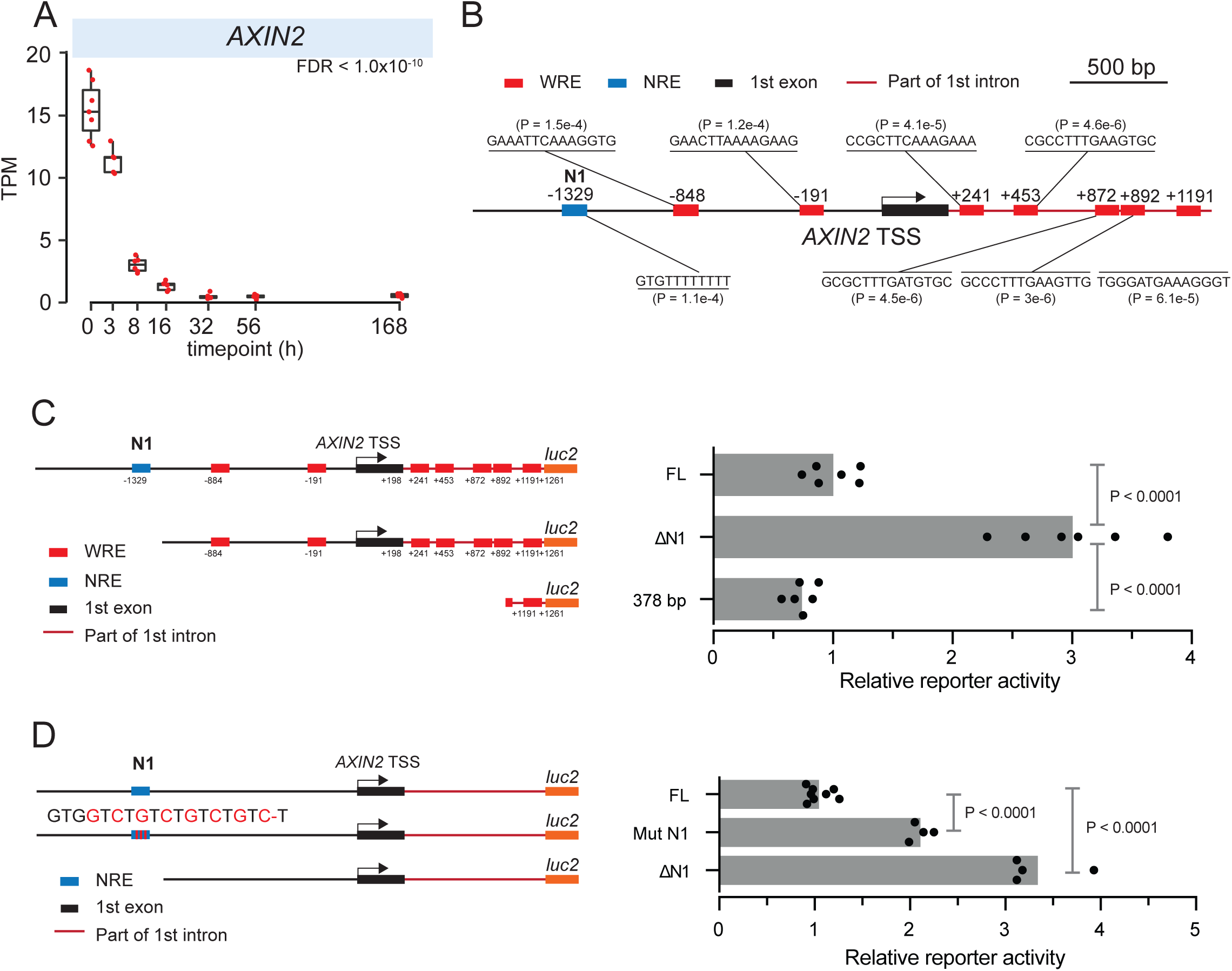
NRE modulates expression of the Wnt-activated/β-catenin-dependent gene *AXIN2*. A) *AXIN2* is a Wnt-activated gene in an orthotopic model of HPAF-II Wnt-addicted cancer where its expression decreases following PORCN inhibition (data from (*7*). B) Positions of putative WREs and NREs in the promoter of *AXIN2*. C) Transcriptional activity of *AXIN2* reporter construct in HCT116 cells following removal of the sequence containing NRE (ΔN1) leads to increased reporter expression. D) Removal or mutation of the NRE increases the transcriptional activity of the *AXIN2* promoter in HCT116 cells.

We confirmed that *AXIN2* is a Wnt-activated/β-catenin-dependent gene in various *in vitro* models (Figure S3). As expected, *AXIN2* was upregulated following BIO treatment in HT1080 cells and was downregulated by using either CRISPRi or siRNA knockdown of β-catenin in HPAF-II and HCT116 cells. In addition, *AXIN2* was significantly upregulated in β-cat4A HPAF-II, and did not respond to PORCN inhibition unlike the HPAF-II WT cells (**Figure S3D**).

To study the functional importance of NREs in the regulation of *AXIN2* expression, we cloned the *AXIN2* promoter region from 2012 bp upstream to 1261 bp downstream of its TSS into a luciferase reporter. Removing the DNA region containing N1 (ΔN1) in this 3273 bp promoter fragment increased the reporter activity by 3-fold in HCT116 cells (**Figure 6C**). Similarly, mutating multiple nucleotides in the poly(thymine) region of NRE N1 to guanine (Mut N1), also significantly enhanced the reporter activity by 2.1-fold (**Figure 6D**). As expected, removing the DNA sequence containing WREs led to a significant decrease in reporter activity (**Figure 6C**). Thus, the NRE can modulate expression of both Wnt/β-catenin repressed and Wnt /β-catenin-activated target genes.

## Discussion

Wnt signaling is a potent regulator of gene expression, which is achieved primarily via changes in nuclear β-catenin abundance. In addition, diverse β-catenin-independent Wnt-regulated (non-canonical) pathways have also been described. However, the contribution of these pathways to the Wnt-regulated transcriptional response is poorly understood. Here, using a PORCN inhibitor that blocks the secretion of all Wnts in a robust orthotopic xenograft cancer model, we find that the majority of Wnt-regulated genes (∼90%) are regulated by changes in β-catenin abundance, indicating that the Wnt-regulated non-canonical pathways, at least in this cancer model, have a small transcriptional impact. Furthermore, this dataset was also interrogated to better understand how Wnt/β-catenin signaling can repress gene expression. This analysis confirms and refines the role of a specific negative Regulatory Element (NRE), extending our understanding of a how β-catenin can repress and/or modulate Wnt-regulated genes.

β-catenin independent roles of Wnt signaling have been well-described. Non-canonical Wnt signaling calcium transients can activate PKC and/or CAMKII, while planar cell polarity signaling functions in part via monomeric GTPase and activation of JNK (*45*, *46*). The consequence of regulating these pathways is cytoskeletal or synaptic reorganization (*47*, *48*). While Wnt-JNK signaling can activate gene expression in Xenopus there is little evidence this pathway regulates transcription in mammals (*49*). Finally, Wnt/STOP signaling increases the proteolysis of proteins including transcription factors such as MYC (*5*, *7*, *50*) but how much they alter gene expression in a physiologic setting is not known. While we cannot separate out the contribution of each of these preceding pathways, our data suggests that taken together, the contribution of these pathways to transcriptional regulation is limited to only 10% of the Wnt-regulated genes. These β-catenin independent genes were enriched for developmental pathways and consistent with that, they showed an enrichment for the transcription factor binding sites for homeobox factors including GSC2, POU6F2 and MSGN1.

There are three strengths of our experimental system. First, we used a cancer model driven by an RNF43 mutation that sensitizes tumor cells to all Wnts, both canonical and non-canonical. Second, using an orthotopic xenograft mode provides a more physiologic milieu, making it far more robust in identifying Wnt-regulated genes than either non-orthotopic xenografts (usually flank) or tissue culture models (*7*). Finally, using a drug that rapidly inactivates Wnt secretion and harvesting the tumors at early time points maximizes the identification of direct targets of the Wnt pathway. This approach provided clear insights into the Wnt-regulated transcriptome.

One striking finding in this and our prior studies is that although there are similar number of genes that are repressed versus activated by Wnt signaling, only a limited number Wnt repressed genes have been identified previously. Here we show that most of these Wnt-repressed genes are still regulated by β-catenin. Several mechanisms have been proposed to explain direct β-catenin/TCF dependent gene repression. This repression was shown to be mediated by binding of TCFs to Wnt Response Elements (WREs). For example, the positioning of WREs in relation to the transcription start site of *MMP7* was shown to be critical for determining its effect on gene expression (*27*). In Drosophila, Wnt/β-catenin signaling represses *stripe* expression at the parasegment boundary during development by steric competition between TCF/LEF (Pangolin) and the transcriptional activator Ci at partially overlapping binding sites (*51*). In mice during hair follicle bud development Wnt/β-catenin signaling represses E-cadherin (*52*) by TCF binding to WRE, which then recruits the transcriptional repressor Snail.

Here we confirm and extend the identification of a TCF-dependent negative regulatory element (NRE), where β-catenin interacts with TCF to repress gene expression. This is consistent with prior work that identified non-canonical TCF binding sites involved in gene repression such as a non-canonical TCF site represses *Ugt36Bc* expression in Drosophila (*28*) and a novel bipartite TCF binding sequence for repression was identified in the fly lymph gland (*29*). Some studies have also suggested that Wnt-mediated repression works by TCF forming a complex with another TF such as GATA3, forming a repressive complex (*41*).

Kim et al. identified an 11 bp repressive motif, termed the Negative Regulatory Element often present alongside the canonical WRE in Wnt-regulated genes (*34*) and was shown to interact with both TCF and β-catenin proteins. Our study confirms that the NRE, albeit with a modified sequence, is enriched in a subset of Wnt-repressed genes. Our mutagenesis studies show that the NRE directs β-catenin dependent repression of the long non-coding RNA *ABHD11-AS1*, and interestingly, also modulates the expression of the robustly Wnt-activated gene *AXIN2*. Overall, this study supports the role of the NRE as an important *cis*-regulatory motif regulating Wnt target genes in human cells and suggests a *cis-*regulatory grammar which can determine whether a target gene will be repressed or activated by Wnt signaling.

We observed 10% of Wnt-regulated genes to be β-catenin independent. This may be due to the high sensitivity of a cancer cell line used in this study that may have shaped the Wnt-regulated transcriptome to favor expression of β-catenin dependent genes. It is also possible that the β-catenin independent gene expression may be more common in normal tissue homeostasis and/or developmental stages. These questions can be addressed in future studies.

In conclusion, this study provides a comprehensive analysis of the β-catenin dependent vs independent genes in a cancer model and advances our knowledge of the role of a cis-regulatory motif in regulating the expression of Wnt target genes in human cells.

## Acknowledgements

This work was supported by National University of Singapore and Yale-NUS College (through Reimagine Research Grant IG20-RRSG-001 to N.H.), and by STAR Award MOH-000155 to D.M.V. from the National Research Foundation Singapore, administered by the Singapore Ministry of Health’s National Medical Research Council. Components of the Bioinformatics analysis was performed on the Cardiff School of Biosciences’ Biocomputing Hub HPC/Cloud infrastructure (Resources funded by the Cardiff School of Biosciences).

## Data availability

RNA-Seq data is available in the NCBI Gene Expression Omnibus (GEO; http://www.ncbi.nlm.nih.gov/geo/) under accession number GSE291732. All analysis code is available at: https://github.com/harmstonlab/wnt_nre_manuscript

## Methods

### Study approval

NOD SCID gamma mice were purchased from InVivos, Singapore and from Jackson Laboratories, Bar Harbor, Maine. All animal studies were approved by the SingHealth Institutional Animal Care and Use Committee and adhered to relevant regulations. The animals were housed in standard cages and had unrestricted access to food and water.

### RNA-seq

Briefly, HPAF-II cells with stable expression of firefly luciferase with and without stable expression of mutant β-catenin were orthotopically injected into the pancreas of NSG mice, as previously described. Approximately 4 weeks later, mice were treated with ETC159 or vehicle as indicated and then sacrificed at the indicated time points. RNAseq was performed on harvested tumors as previously described (*7*).

### Data processing and quality control

Xenome was used to remove murine (mm10) reads (*53*) and FastQC to ensure the overall quality of the sequences. The remaining reads were then aligned against hg38 (Ensembl version 100) using STAR v2.7.1a (*54*) and RSEM v1.3.1(*55*). Genes which had less than 10 reads mapping on average over all samples, as well as reads mapping to rRNA, mtRNA, snoRNA, and snRNA were filtered out. Differentially expressed genes were identified using DEseq2 (*56*). Independent filtering was not used in this analysis.

### Identification of β-catenin-dependent and independent clusters

DESeq2 was used to identify genes that responded differently to ETC-159 treatment depending on CTNNB1 status. Gene expression changes were modelled as y ∼ condition + timepoint + condition:timepoint, where condition is wildtype (WT) or mutant (Mut) and timepoint is 0 h, 4 h, 16 h, 56 h. Likelihood ratio tests were also performed to identify genes that changed expression significantly across time within conditions. Pairwise comparison using Wald test was performed between WT and Mut conditions at 0 h to identify genes with differences in baseline expression. Coefficients from the model (representing log fold changes) were clustered using k-means clustering, with the value of k being determined using the elbow criterion.

### Functional enrichment analysis

Gene Ontology (GO) enrichment was performed using enrichGO and pathway enrichments using enrichKEGG from ClusterProfiler (*57*) using all expressed genes as background. Terms with FDR < 0.1 were defined as being significantly enriched.

### Motif enrichment analysis

Promoters were defined as 2000 bp upstream and 500 bp downstream or as stated in the results. Enrichment analysis was performed using monaLisa (*58*) using JASPAR2020 (*59*), min.score 80%, binomial test, all promoters used as background, genome oversample 20.

### NRE motif

The NRE motif was derived from the sequences reported (*34*), however the sequences for Brachyury-1 and Brachyury-2 were reverse complemented before generation of the position frequency matrix.

### ChIP-seq

CTNNB1 and TCF7L2 ChIP-seq data was downloaded from GEO (*40*, *41*). FastQC was used to perform quality checks on raw sequence data and adapters were trimmed using cutadapt. Reads were aligned against the human genome (hg38) using BWA (*60*) and peaks were identified using MACS2 (*61*). ChIPQC was used to assess the quality of ChIP-seq samples and experiments (*62*). For samples where replicates were available, an irreproducible discovery rate (IDR) threshold of 0.05 was used to select for highly reproducible peaks (Li et al. 2011). Peaks were centered on their midpoint and resized to 500 bp for analysis.

Motif enrichment and identification was performed using MEME (*63*). SpaMo was used to determine whether there were significantly enriched spacings between the NRE and WRE motifs (*64*). Default parameters used with the margin size of 150bp.

### Motif identification

*ABHD11-AS1* promoter region (3.4 kb upstream and 200 bp downstream of its TSS) and *AXIN2* promoter region (3.5 kb upstream to 1 kb downstream of its TSS) were used as input to run FIMO to scan for putative NRE and WRE sites (*65*).

### Construction of ABHD11-AS1 and AXIN2 promoter reporters to assess the effect of NRE truncation and mutation on promoter activity

*ABHD11-AS1* promoter region from 3327 bp upstream to 151 bp downstream of its TSS and *AXIN2* promoter region from 2012 bp upstream to 1261 bp downstream of its TSS were cloned from genomic DNA with primer sequences listed in table S3. The promoter regions were cloned into the luciferase reporter pGL4.20 (Promega) with NheI and HindIII restriction sites (named as FL construct). A series of deletion constructs were generated using FL as a template with primers sequences listed in table S3. All PCR products were digested with NheI and HindIII and cloned into pGL4.20 (Promega). Site-directed mutagenesis was performed to mutate NRE with primers listed in table S3.

### Construction of minimal reporter

311 bp (without any putative NREs or WREs) from the ABHD11-AS1 promoter region (sequences listed in table S3) was cloned into the pGL4.20 (basic vector with no promoter) with NheI and HindIII restriction sites to construct a MimRep. Two NREs and 2 WREs sequences (listed in table S3) were cloned into the MimRep with SacI and KpnI restriction sites to construct 2NRE-Reporter and 2WRE-Reporter.

### Luciferase assay

HT116 or HT1080 cells were seeded into 24-well plates one day before transfection. Cells were transfected with different constructs and control Renilla luciferase expression vector using Lipofectamine 2000 (Invitrogen) according to manufacturer’s instructions. Luciferase activity was assessed 24h after transfection with the Dual-Luciferase Reporter Assay System (Promega). Transfections were performed at least in triplicate on at least two separate experiments. Luciferase signals were first normalized to Renilla. The relative amount of luciferase activity was further normalized to the empty vector (pGL4.20) transfected cells.

### CRISPRi Knock down studies

The sgRNAs were cloned into doxycycline-inducible lentiviral sgRNA expression vector FgH1tUTG as previously described. The sgRNA plasmid was packaged into lentiviral particles with psPAX2 and pMD2.G packaging plasmids. The virus supernatant was harvested 48 and 72 h after transfection, filtered through 0.45-μm filter, and stored at − 80 °C. For individual sgRNA knockdown using doxycycline-inducible lentiviral sgRNA expression vector FgH1tUTG, 1 μg/ml doxycycline final concentration (dox) (from a stock of 10 mg/ml dissolved in DMSO) was used to induce sgRNA expression from the system, while DMSO was used as the control. After 48 h induction, total RNA was isolated from the CRISPRi knockdown cells. RT-qPCR was performed to assess the knockdown efficiency for *CTNNB1* with *EPN1* gene as an internal control. RT-qPCR primers are listed in table S3.

---

**Figure S1.**
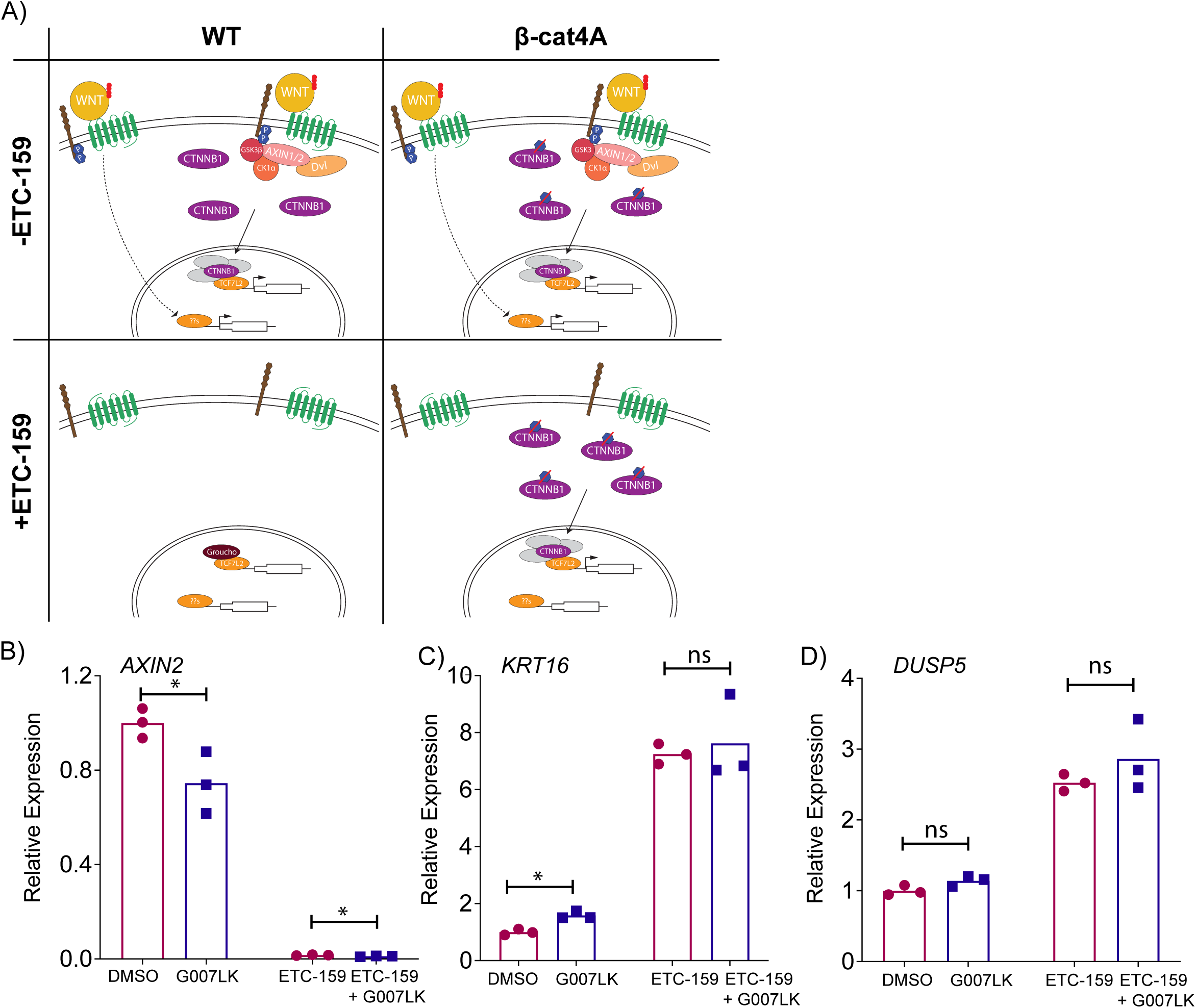
A) Schematic of the effects of PORCN inhibition on the WNT signaling pathway in HPAF-II WT and β-cat4A cells. B) *AXIN2* expression is responsive to tankyrase inhibition while *KRT16* (IC2) and *DUSP5* (IC2) are not.

**Figure S2.**
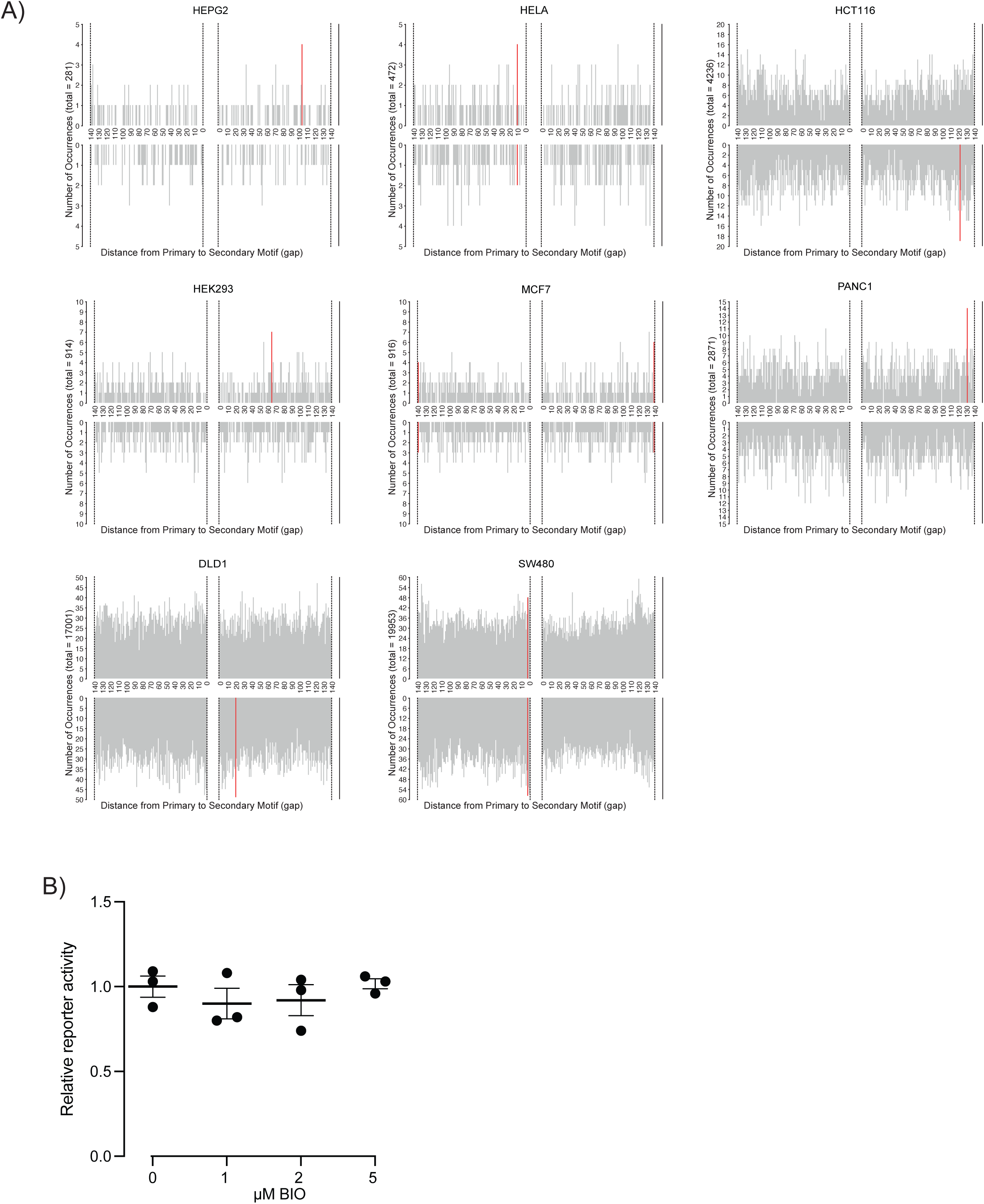
A) Investigating distances between WRE and NRE motifs using SPAMO identified no preferred spacing between the two motifs. B) Non-Wnt responsive reporter response to BIO in HT1080 cells - The reporter is 160(+) to 151(-) ABHD11-AS1 TSS

**Figure S3.**
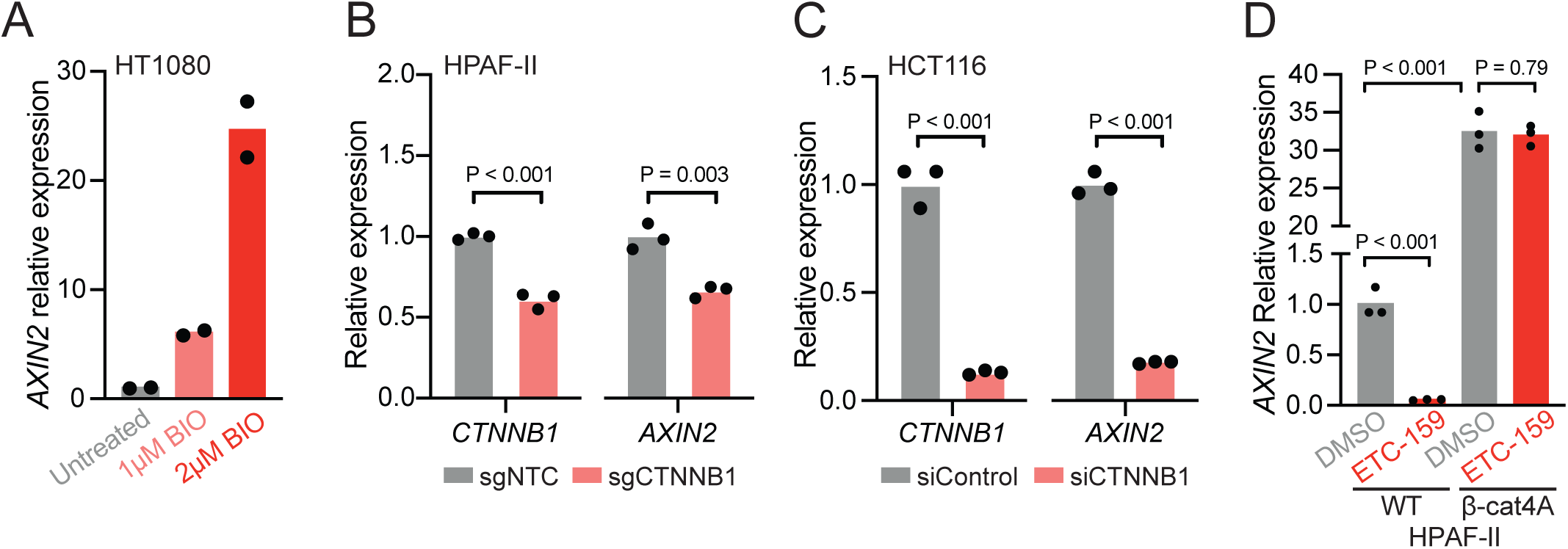
A) AXIN2 expression is increased following β-catenin activation using BIO in HT1080 cells. Inhibiting β-catenin using B) CRISPRi or C) siRNA leads to a decrease in *AXIN2* expression in HPAF-II and HCT116 cells respectively. D) Expression of *AXIN2* decreases in WT but not β-cat4A cells following treatment with ETC-159.

